# Transcriptomic and Proteomic Analysis Reveals Nitrogen Recycling as a Core Mechanism for *Prochlorococcus* Prolonged Survival

**DOI:** 10.1101/2025.11.24.690089

**Authors:** Osnat Weissberg, Dikla Aharonovich, Daniel Sher

## Abstract

*Prochlorococcus*, the dominant cyanobacterium in the oligotrophic ocean, possesses a streamlined genome and depends on interactions with heterotrophic bacteria for survival under various stressors. While the role of ‘helper’ bacteria in mitigating oxidative stress is established, the mechanisms enabling its long-term survival under nitrogen (N) limitation remain poorly characterized.

Here, we employ a multi-omics approach—integrating transcriptomics and proteomics—to investigate the physiological processes that facilitate the prolonged survival of *Prochlorococcus* in co-culture with the marine heterotroph *Alteromonas* during conditions of extreme N-deprivation.

Our results demonstrate that, unlike axenic cultures which rapidly perish, *Prochlorococcus* in co-culture maintains viability for months following the depletion of initial N-sources. Molecular analysis identifies a shift in both organisms that underpins this persistence: *Prochlorococcus* strongly upregulates high-affinity N-scavenging pathways, while *Alteromonas* exhibits transcriptional and translational changes consistent with increased organic matter degradation and reduced motility. This suggests that *Alteromonas* functions as a key nitrogen recycler, providing a continuous, albeit low-level, supply of bioavailable NH_4_^+^ to its photoautotrophic partner through the remineralization of organic matter.

These findings support and extend the Black Queen Hypothesis, illustrating that the benefits conferred by heterotrophs to genome-streamlined primary producers encompass not only the detoxification of reactive oxygen species but also the continuous provisioning of essential macro-nutrients under starvation conditions. This tightly coupled, mutualistic relationship represents a critical factor driving the resilience and productivity of microbial communities in oligotrophic marine ecosystems.

## Introduction

One of the dominant forces in global carbon fixation is the cyanobacterium *Prochlorococcus*. *Prochlorococcus* is abundant in the oligotrophic areas of the world ocean, especially near the bottom of the photic zone. Despite its minute cell size, this bacterium is estimated to contribute 10-20% of global photosynthesis (Flombaum et al., 2013).

*Prochlorococcus* has a small, streamlined genome and has lost many genes (Kettler et al., 2007). Specifically, *Prochlorococcus* lacks catalase-peroxidase (katG), necessitating the presence of a ‘helper’ heterotrophic bacterium to mitigate oxidative stress (Morris et al., 2012). This observation supports the Black Queen Hypothesis, which postulates that the organism’s survival despite core gene loss is enabled by co-occurring bacteria providing the requisite enzymatic products as a public good (Morris et al., 2012).

Studies of *Prochlorococcus* interactions with marine heterotroph bacteria focused on the marine copiotroph, *Alteromonas*. This heterotroph facilitates *Prochlorococcus* survival under multiple stressors. *Prochlorococcus* was shown to benefit from the heterotroph presence under multiple conditions including extended temperature range and increased oxidative stress (Ma et al., 2018), long term nitrogen limitation (Weissberg et al., 2023), extended darkness (Biller et al., 2018; Coe et al., 2024), and competition from other phytoplankton (Calfee et al., 2022).

Our investigation specifically addressed the capacity of *Alteromonas* to promote the long-term survival of *Prochlorococcus* under nitrogen-limited conditions (Weissberg et al., 2023). We utilized proteomic and transcriptomic sequencing to elucidate the physiological mechanisms governing this positive interaction.

Our study focused on nitrogen limitation; this is a common condition in the oligotrophic gyres (Moore et al., 2013). *Prochlorococcus* response to nitrogen starvation and nitrogen limitation includes upregulation of enzymes involved in nitrogen metabolism and nitrogen transporters, including ntcA, the nitrogen regulator in *Prochlorococcus (Domínguez-Martín et al., 2017; Tolonen et al., 2006)*. Additionally, photosynthesis slows down and a larger portion of fixed carbon is released into the environment (Szul et al., 2019; Tolonen et al., 2006). Finally, translation is slowed down coupled with a reduction in ribosomal activity, hypothesized to be related to the conservation of nitrogen, as ribosomes are nitrogen rich (Domínguez-Martín et al., 2017).

Although *Prochlorococcus* has been extensively studied under nitrogen limitation and in heterotrophic interactions, the specific mechanisms underlying its long-term survival under nitrogen deprivation and the associated helper function remain uncharacterized. We grew *Prochlorococcus* and *Alteromonas* (in coculture and separately) for 90 days and studied their response to nutrient limitation and their interactions. The co-cultivation established a mutualistic interaction, resulting in a protracted period of joint long-term survival. Conversely, the axenic *Prochlorococcus* cultures exhibited rapid decline and loss of viability, while axenic *Alteromonas* concentrations stabilized at an order of magnitude lower population density. We therefore used transcriptome and proteome assays to compare the physiological state in coculture to the axenic controls with the goal of uncovering the underlying interaction mechanisms.

## Results

### Mutualistic interaction between *Prochlorococcus* and *Alteromonas*

*Prochlorococcus* and *Alteromonas* were grown in coculture and separately (axenically) in PRO99-lowN for 90 days, encompassing exponential growth, decline, and (in co-culture) long term survival together. In this medium, *Prochlorococcus* becomes nitrogen-limited following the exponential growth stage (i.e. decline and long-term starvation) (Grossowicz et al., 2017), while *Alteromonas* is expected to be carbon-limited (Weissberg et al., 2025).

When grown in coculture, both strains benefitted from the presence of the other exhibiting mutualistic interaction. The synergistic effects of the interaction are maximally evident during the decline and long-term starvation phases. *Prochlorococcus* grew exponentially and its growth rates were the same in the coculture and the axenic control. Upon depletion of NH4+, the differences between the treatments begin to emerge. Axenic *Prochlorococcus* exhibited a rapid decline in cell number and loss of viability (Fig 1B), whereas the co-cultured *Prochlorococcus* entered a protracted, slow decline that persisted until the end of the experiment (fig 1A). This is corroborated by viability assays: axenic *Prochlorococcus* does not revive when transferred into fresh media on day 11. Meanwhile, the cocultures *Prochlorococcus* in the coculture is viable on days 30 and 60 and becomes unviable only at the end of the experiment on day 89 (sup fig S1B).

**Fig 1:**
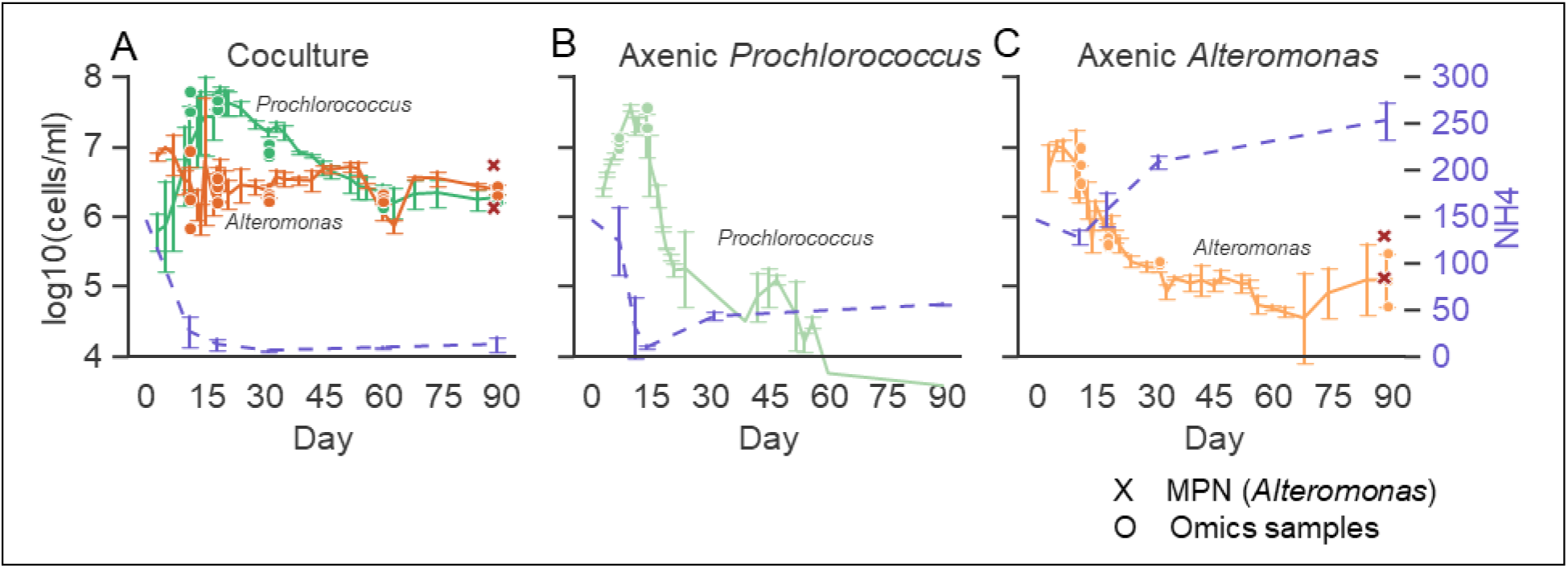
growth of *Prochlorococcus* and *Alteromonas* A) in coculture B) Axenic *Prochlorococcus*. C) Axenic *Alteromonas* as measured by flow cytometry. Large dots indicate samples for omics (RNASEQ/proteome), large X in the axenic *Alteromonas* and the co-cultures represent culturable heterotroph counts (MPN).

*Alteromonas* go into decline at the beginning of both axenic growth and co-culture. In coculture, this decline lasts until NH_4_^+^ is depleted and *Prochlorococcus* starts to decline. At this point, *Alteromonas* slowly recovers, till ∼day 50, where we observe a second, shorter decline and recovery (fig 1A). In the Axenic *Alteromonas* cultures, the decline slows down around day 30 but continues till the end of the experiment (fig 1C). *Alteromonas* cell numbers are 1-2 orders of magnitude higher in coculture (2 * 10^6^ *Alteromonas* cell ml^-1^ in coculture vs 10^5^ cell ml^-1^ in axenic control (measured as mean of the flow cytometry measurements on days 60-90). Both treatments of *Alteromonas* are viable at the end of the experiment as measured by serial dilutions in marine broth (MPN, fig 1). The number of viable *Alteromonas* cells in coculture is an order of magnitude higher than those of the axenic *Alteromonas* (41±24 10^5^ cell ml^-1^ vs. 3±2 10^5^ cell ml^-1^ respectively).

Cell counts for *Alteromonas* oscillate. These oscillations in the coculture are more pronounced and are coupled with smaller oscillations in *Prochlorococcus* growth. There are several possible explanations for these dynamics. One possible explanation involves recycling of dead cells. When *Prochlorococcus* population collapses, a lot of nutrients are released, which supports *Alteromonas* growth. The expanding *Alteromonas* population remineralizes a fraction of these nutrients, converting them into forms bioavailable for *Prochlorococcus*, particularly nitrogen compounds like ammonium and, potentially, amino acids (see below), thereby establishing a cyclical process

Other options include evolutionary processes, such as the GASP phenotype in *E. coli* (Finkel, 2006), where the observed cycles are caused by emergence of better adapted subpopulations. We currently do not have genome sequences that could help us differentiate between these two options.

These growth and death dynamics are coupled with NH_4_^+^ concentrations. In the coculture, NH_4_^+^ level steadily declines while *Prochlorococcus* is in exponential growth. NH_4_^+^ levels remain low throughout the rest of the experiment (fig 1A). These persistently low values likely indicate rapid ammonium turnover, wherein any newly produced NH4^+^ is quickly assimilated by the microbial cells (Klawonn et al., 2019). In axenic *Alteromonas*, NH_4_^+^ levels go up and are higher than the initial levels in the media, indicating active ammonium production via enzymatic activity or as result of cells breaking down. Ammonium levels also go up, though to a lesser extent, in axenic *Prochlorococcus* cultures. We hypothesize this accumulation results from the decomposition of senescent cells, given that *Prochlorococcus* is not viable at these timepoints.

### *Prochlorococcus* shifts from exponential growth to survival or death

We used transcriptome and proteome analyses to assess the physiological state of the *Prochlorococcus* and *Alteromonas* cells as they each grew alone, and together. Our sampling points are mostly during long-term decline as that is the focus of our study (beginning of decline on day 14 and long-term on days 30, 60, 90). These were compared to exponential growth (for *Prochlorococcus*, days 7/11). Finally, we used axenic controls (fig 1).

In both the omics assays, samples cluster by treatment and sampling time with a clear difference between early and late timepoints, although we note that, for the proteome samples, this could be due to the different databases used.

Based on the number of differentially expressed genes and differentially abundant proteins, *Prochlorococcus* growth and death can be separated into clear phases. In the Axenic controls, there were many up- and down-regulated genes when comparing exponential growth and decline (figure 2A). The late timepoints (30 and 90 days) likely represent decomposing dead cells (we could not obtain RNA from these samples, and the proteome data is likely mostly from cell debris). The much slower decline of *Prochlorococcus* in co-culture is mirrored by a gene and protein expression profile that is, overall, much more similar to the exponential growth than to the axenic death (Figure 2B). These patterns were also seen in the Principal Component Analysis, where major differences were seen between growth and death in the axenic controls (along the X axis, which explains most of the diversity), whereas the exponential growth and decline in co-culture, while clearly different, were closer to each other (Figure 2C, D). Interestingly, the gene expression of the sample taken after 30 days in coculture clustered with the earlier decline stage (day 18), whereas the proteome clustered with the later, long-term survival stages (days 60,90). This may suggest this time-point as a midpoint in the physiological shift the cells are undergoing.

**Fig 2:**
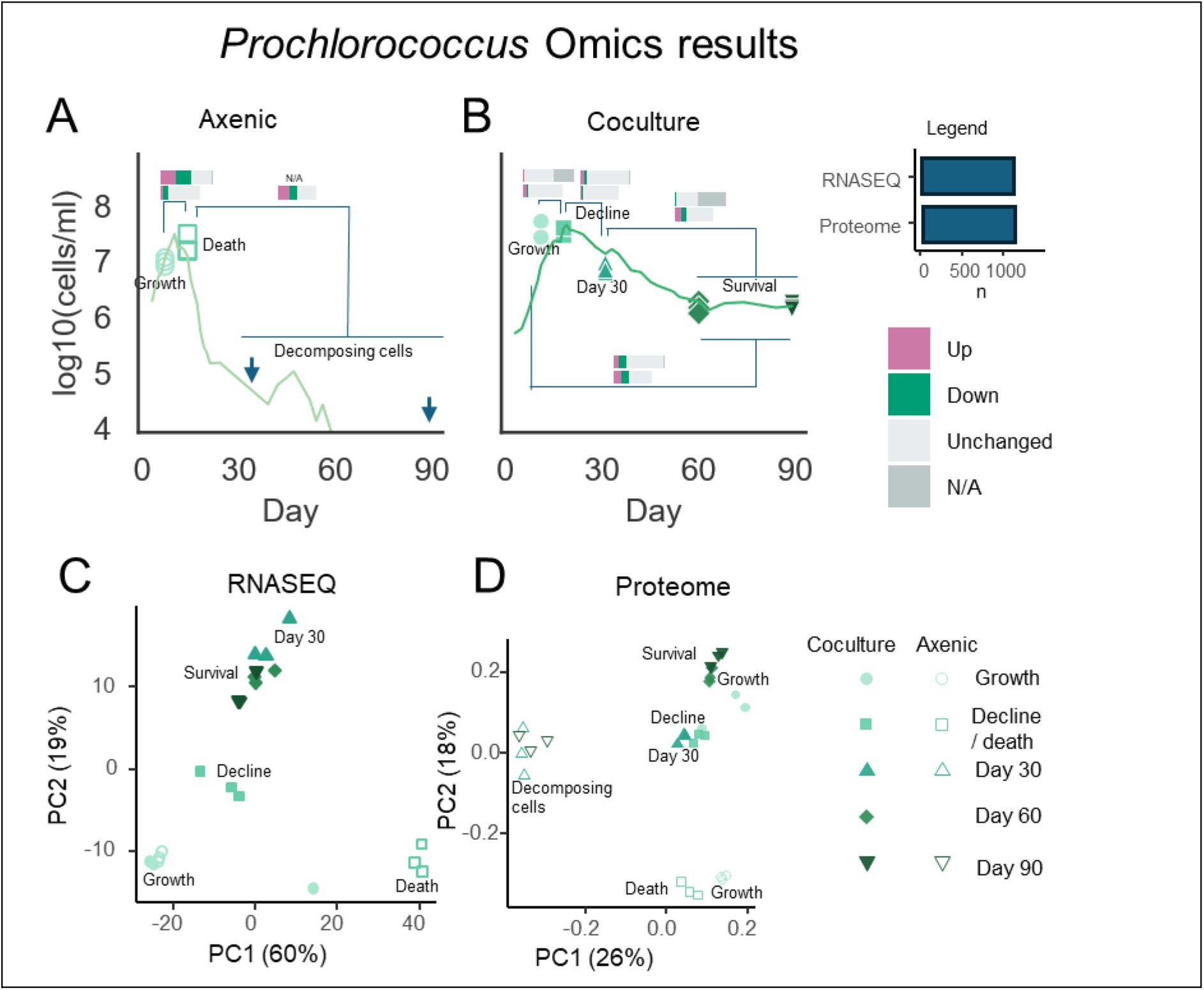
Omics results for *Prochlorococcus*. A-B): number of differentially expressed genes in comparisons between different omics timepoints for Axenic *Prochlorococcus* (A) and *Prochlorococcus* in coculture (B). C-D) Principal Component Analysis of *Prochlorococcus* gene expression (C) and Proteome (D). In panels A-B omics timepoints are highlighted by large markers or arrows. Arrows represent timepoints where cell concentration is below limit of detections. Differential expression is N/A when read count is too low or for outliers flagged by DESeq2.

It is also noteworthy that the gene expression during exponential growth was similar between the axenic controls and co-cultures, suggesting a minimal effect of the interaction with the heterotrophs at this stage.

### Differentially expressed genes and proteins in *Prochlorococcus*

How do Prochlorococcus cells surviving with Alteromonas for extended periods of N starvation differ from those growing exponentially? We assessed the physiological state of *Prochlorococcus* by comparing measurements taken during exponential growth to later timepoints. For the cocultures, we compared measurements taken late in the experiment (long-term decline, days 60, 90) to exponential growth (day 11). The axenic culture died around day 11, therefore we compared dying cultures (day 11) to exponential growth (day 7).

There is major downregulation of gene expression when the cells are stressed and likely dying, with a strong signal of photosynthesis genes. A similar bias towards down-regulation is seen also in the proteome, although at a lower magnitude, and a dominance of general metabolic genes. In contrast, changes in gene expression and protein abundance are much more “balanced” when comparing growth and survival (fig 3).

**Fig 3:**
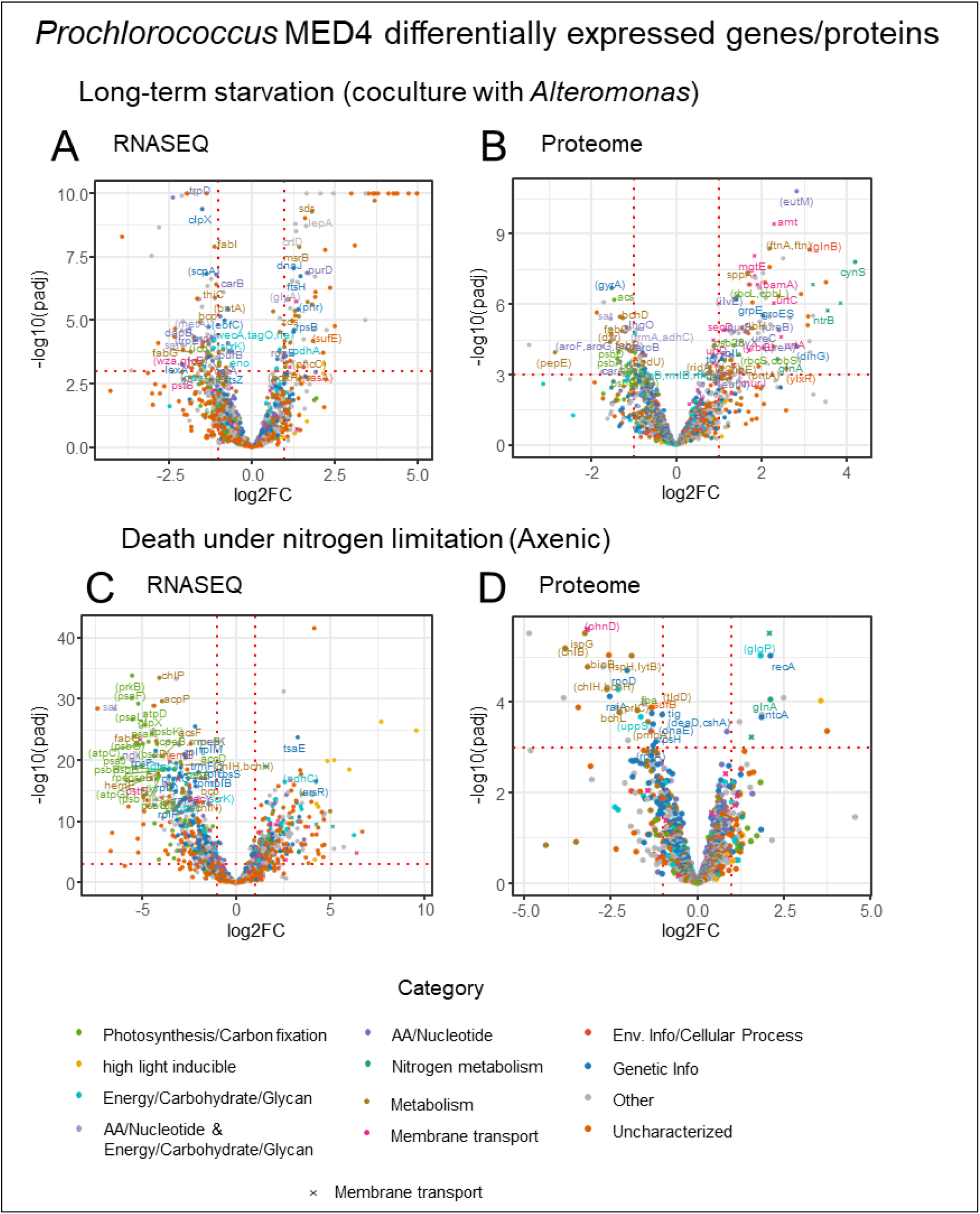
differentially expressed genes in *Prochlorococcus* long-term decline in co-culture with *Alteromonas*. A. differentially expressed genes (RNASEQ). B. Differentially abundant proteins (Proteome). And in axenic death stage (C: RNASEQ, D: Proteome). Colored by the most differentially expressed categories. X signs represent membrane transporters.

In addition, *Prochlorococcus* responded by slowing down photosynthesis and upregulating nitrogen metabolism (fig 3). High-light inducible proteins were upregulated, these genes are involved in protecting the photosynthetic mechanism during stress response.

In the proteome, there are multiple upregulated membrane transporters. And there is a strong response (both upregulation and downregulation) of genes related to genetic information processing and to metabolism of amino acids and nucleotides, both nitrogen-containing compounds.

Many of the genes that are differentially expressed in the *Prochlorococcus* transcriptome are uncharacterized proteins. In fact, all but one of the top 5 upregulated and downregulated transcripts are uncharacterized (Table 1).

**Table 1:** Top *Prochlorococcus* upregulated genes and proteins in long-term survival.

| Gene id | gene | product | KEGG pathway | Proteome |  | RNA |  |
| --- | --- | --- | --- | --- | --- | --- | --- |
|  |  |  |  | logFC | Padj | logFC | Padj |
| TX50_RS09500 |  | hypothetical protein | NA | NA | NA | 5.29 | 3E-26 |
| TX50_RS04605 |  | hypothetical protein | NA | NA | NA | 5.08 | 8E-20 |
| TX50_RS09690 |  | hypothetical protein | NA | NA | NA | 4.75 | 1E-18 |
| TX50_RS09290 |  | hypothetical protein | NA | NA | NA | 4.41 | 5E-18 |
| TX50_RS01960 |  | hypothetical protein | NA | NA | NA | 4.38 | 2E-21 |
| TX50_RS01985 | cynS | cyanase | Nitrogen metabolism | 4.22 | 2E-08 | 1.10 | 0.03 |
| TX50_RS02045 |  | SIMPL domain-containing protein | Function unknown | 3.53 | 1E-07 | 1.05 | 0.16 |
| TX50_RS01975 | ntrB | nitrate ABC transporter permease | Nitrogen metabolism; ABC transporters | 3.57 | 2E-06 | 0.56 | 0.53 |
| TX50_RS01970 |  | ABC transporter substrate-binding protein | Nitrogen metabolism; ABC transporters | 3.88 | 9E-07 | 0.34 | 0.77 |
| TX50_RS02565 |  | PepSY domain-containing protein | NA | 3.52 | 0.01 | -0.28 | 0.48 |

In the proteome, all the top differentially expressed proteins are also differentially expressed in the transcriptome, though to a lower extent. 3 of the top upregulated proteins are part of nitrogen metabolism, which is reasonable when considering that *Prochlorococcus* is nitrogen limited in the experiment media (Grossowicz et al., 2017). The five top down-regulated proteins include enzymes involved in glycan metabolism and a peptidase (Table 2).

**Table 2:** Top *Prochlorococcus* downregulated genes and proteins in long-term survival.

| Gene id | gene | Product | KEGG pathway | Proteome |  | RNA |  |
| --- | --- | --- | --- | --- | --- | --- | --- |
|  |  |  |  | logFC | Padj | logFC | Padj |
| TX50_RS09770 |  | hypothetical protein | NA | NA | NA | -4.29 | 6E-03 |
| TX50_RS09810 |  | hypothetical protein | NA | NA | NA | -3.93 | 5E-09 |
| TX50_RS00165 |  | transcription factor TFIID | NA | NA | NA | -3.65 | 1E-03 |
| TX50_RS07860 |  | hypothetical protein | NA | NA | NA | -3.15 | 1E-03 |
| TX50_RS07415 |  | LOG family protein | NA | -1.54 | 0.05 | -3.04 | 3E-08 |
| TX50_RS05775 |  | oligoketide cyclase | NA | -3.45 | 5E-05 | -1.74 | 1E-12 |
| TX50_RS06610 | (wecA, tagO, rfe) | undecaprenyl/decaprenyl-phosphate alpha-N-acetylglucosaminyl 1-phosphate transferase | Glycan biosynthesis and metabolism | -3.12 | 2E-03 | -1.18 | 7E-05 |
| TX50_RS07345 |  | GAF domain-containing protein | NA | -2.49 | 2E-03 | -0.89 | 0.01 |
| TX50_RS05300 | (pepE) | peptidase E | Peptidases and inhibitors | -2.84 | 1E-04 | 0.09 | 0.89 |
| TX50_RS03935 |  | hypothetical protein | NA | -1.91 | 2E-03 | 0.39 | 0.36 |

### *Prochlorococcus* physiological state in survival and death modes

What is the physiological state *Prochlorococcus* in the coculture? We used enrichment analysis of the KEGG pathways to get a system level perspective. The major physiological changes include increased expression of genes and abundance of proteins involved in nitrogen metabolism (fig 4). As observed with individual genes, metabolism seems to slow down (including translation machinery and photosynthesis), as suggested by the overall down-regulation of the pathways for Ribosome, Translation and genetic information processing. Both responses were also observed in experiments where *Prochlorococcus* was nitrogen limited or nitrogen starved (Felcmanová et al., 2017; Szul et al., 2019; Tolonen et al., 2006). In addition, membrane transport is upregulated in the survival phase in the proteome. There are very few differentially expressed pathways in survival transcriptome as most of the differentially expressed genes have not been characterized (are hypothetical).

**Fig 4:**
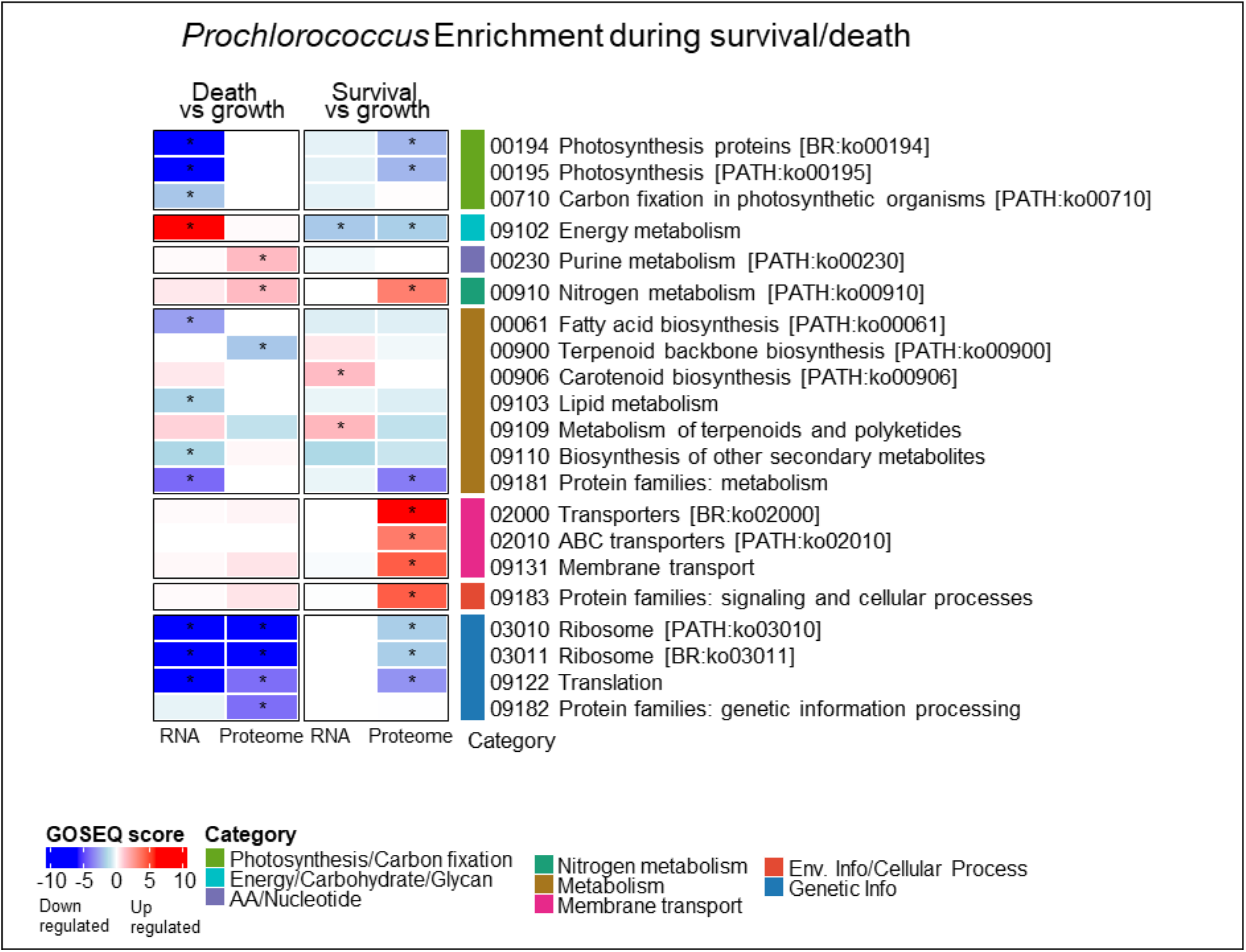
Enriched KEGG pathways in *Prochlorococcus* during Axenic death and long-tern survival in coculture. Pathways enriched when comparing death and long-term survival (days 60,90) to log exponential.

### Decomposing *Prochlorococcus* cells

Axenic *Prochlorococcus* was sequenced on 4 timepoints: growth (days 7), decline (day 11) and dead cells (days 30 and 90). On the 2 latter points, only proteome data is available as there was insufficient RNA for extraction. At these points, *Prochlorococcus* has not been viable for weeks, and the sample contains either vital but not viable “zombie cells” (i.e. that cannot reproduce but are still metabolically active, (Roth-Rosenberg et al., 2020), dying ones or decomposing cell debris. One way to interpret this is as proteins with long lifetime. During the decomposition stage, there is decreased abundance of proteins involved in general metabolism, photosynthesis and membrane transport. In contrast there is increased relative abundance of proteins involved on genetic information processing (Ribosomes, RNA biosynthesis and degradation, transcription and translation), in the exosome and cell death, and in metabolism of amino acids and nucleotides.

**Fig 5:**
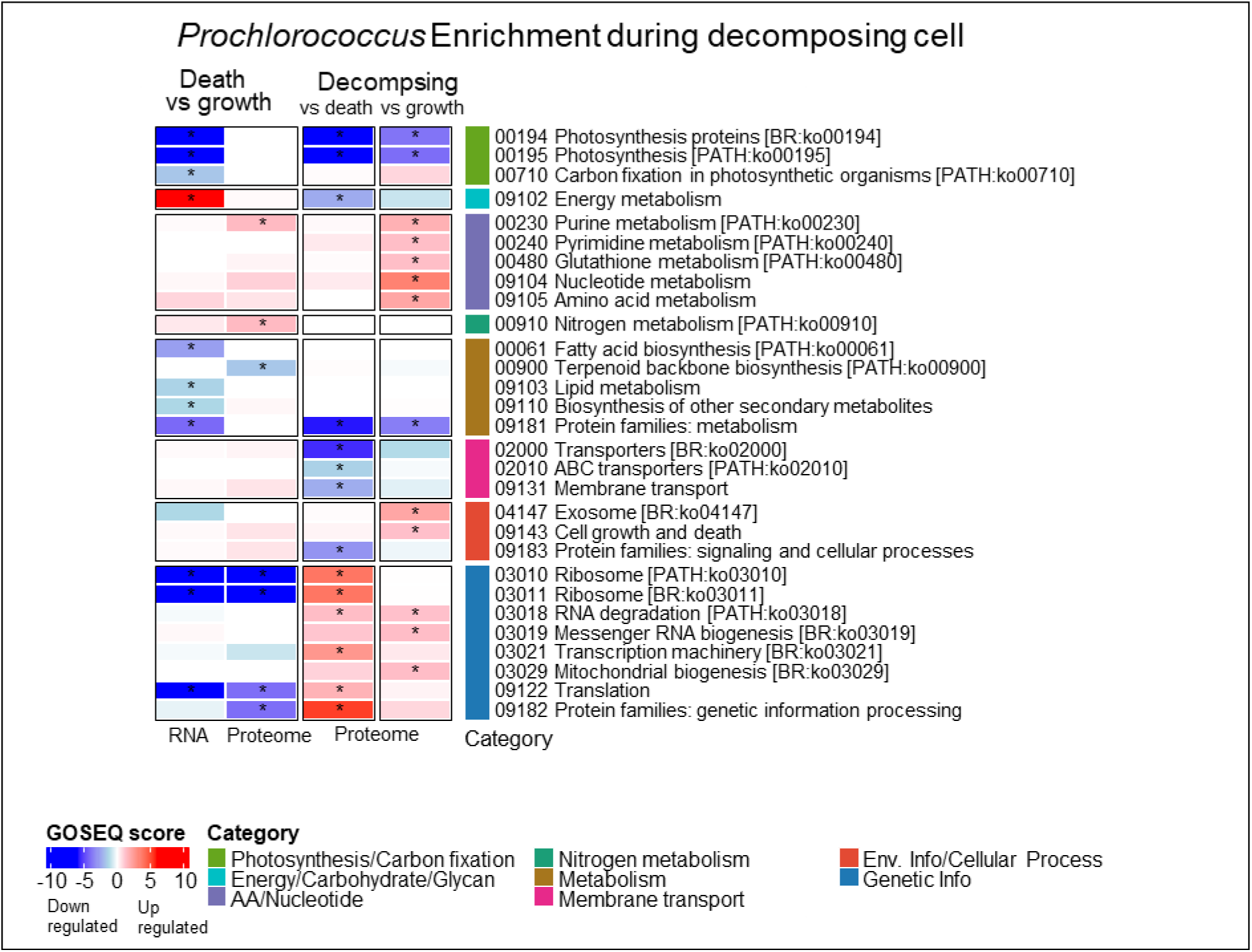
KEGG enriched pathways in the axenic *Prochlorococcus* cultures and in decomposing *Prochlorococcus* cell (proteome on days 30, 90).

### *Alteromonas* response is continuous

*Alteromonas* response is different from that of *Prochlorococcus*. The experimental media is not favorable for *Alteromonas*, as we did not add any organic carbon source. There is no exponential growth stage. Instead, the cell numbers begin to decline as the experiment starts, likely due to C starvation. In the axenic cultures this decline progresses until the cell level stabilizes around 10^5^ cell ml^-1^(fig 6A). In the coculture, *Alteromonas* recover around day 15 (concordant with NH_4_^+^ depletion and *Prochlorococcus* decline) and relatively stable population of ∼10^6^ cell ml^-1^ (fig 6B).

The omics results reflect these dynamics. In the transcriptome, there is a clear metabolic shift as the culture ages, this is reflected by all samples being on a line in the PCA. Interestingly, the cocultures exhibit an accelerated reaction, each sample resembling the previous time point of the axenic controls (coculture day 11 resembles axenic day 18, and coculture day 18 resembles axenic day 30 transcriptions). The transcriptome of the late cocultures (days 60,90) is remarkably different than that of all previous samples.

The proteomes cluster by age. In the axenic *Alteromonas* there is a similar progression while in the coculture, samples of days 30, 60, 90 cluster aways from the earlier samples.

**Fig 6:**
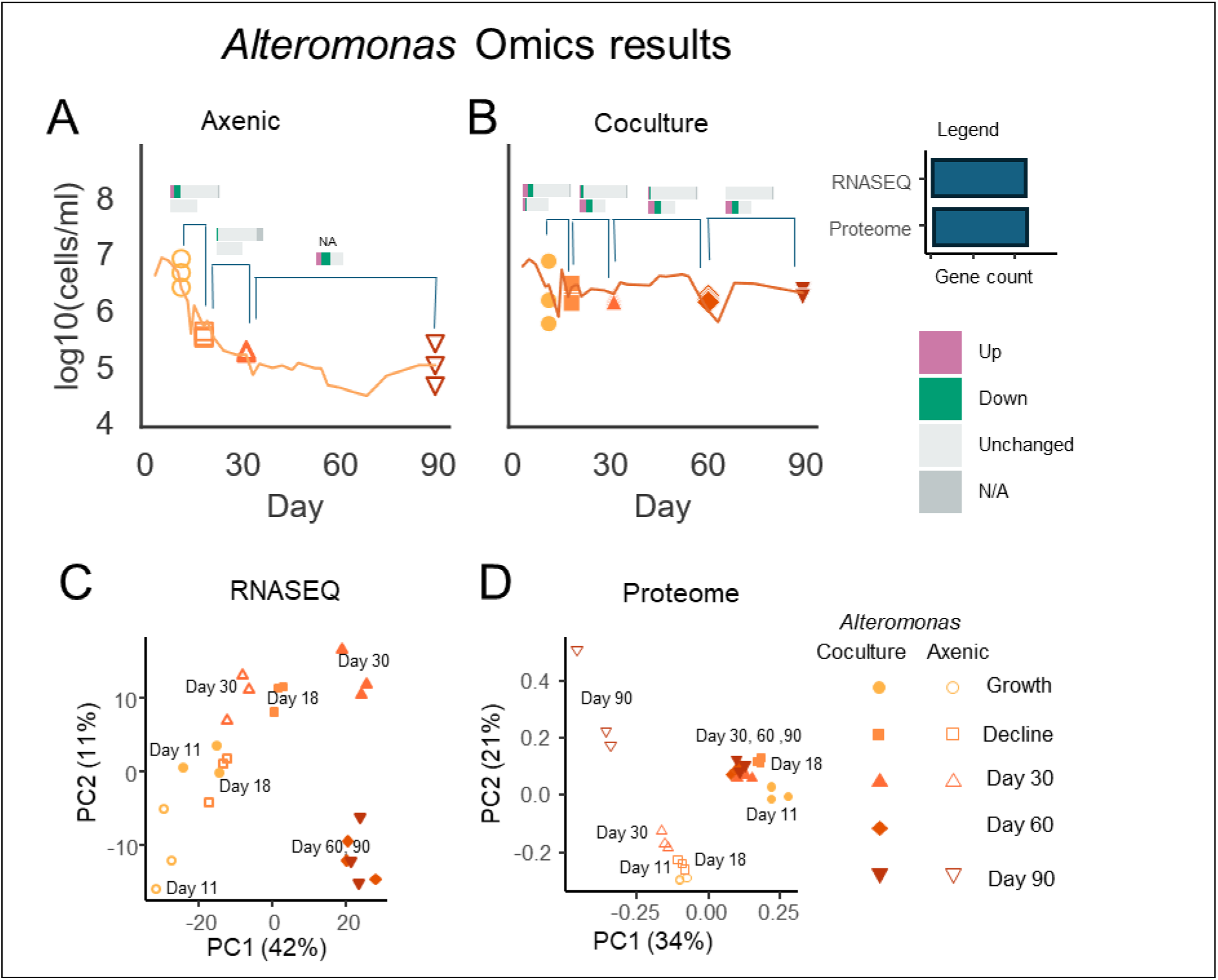
Omics results for *Alteromonas*. A-B: differentially expressed genes in different omics timepoints for A) Axenic *Alteromonas* B) *Alteromonas* in coculture. C) *Alteromonas* RNASEQ. C. *Alteromonas* Proteome. In panels A-B omics timepoints are highlighted by large markers or arrows. Arrows represent timepoints where cell concentration is below limit of detections. Differential expression is N/A when read count is too low or for outliers flagged by DESeq2.

### Differentially expressed genes and proteins in *Alteromonas*

Zooming in on the later timepoints (60,90) where *Alteromonas* is in mutual interaction with *Prochlorococcus*, there is a clear response. Many of the genes that are differentially expressed relative to the early timepoints are related to energy, metabolisms of amino acids and nucleotides, membrane transport and motility. Nitrogen metabolism is upregulated in the proteome.

**Fig 7:**
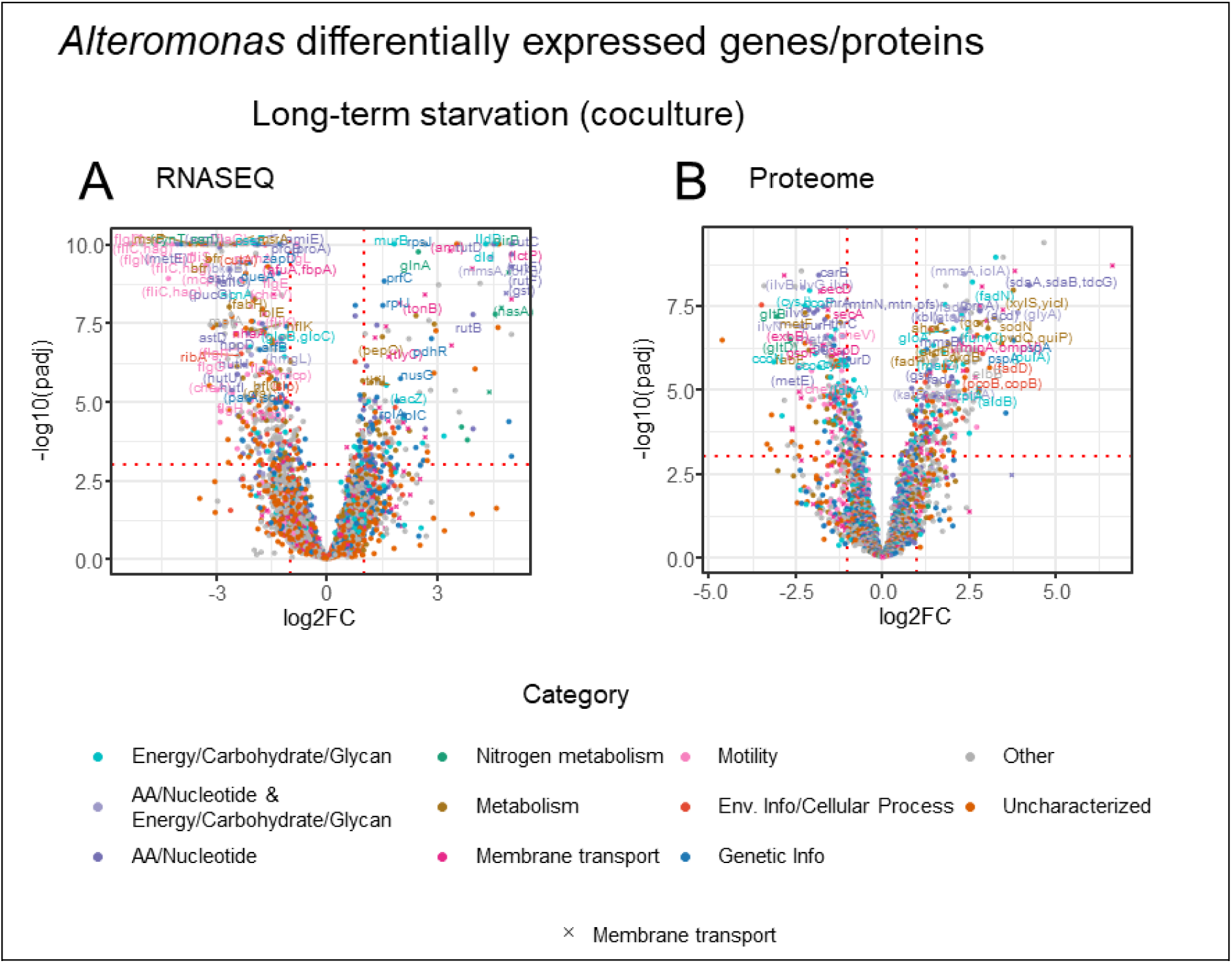
differentially expressed genes in *Alteromonas* long-term survival in co-culture. A. differentially expressed genes (RNASEQ). B. Differentially abundant proteins (Proteome). Colored by the most differentially expressed categories. X signs represent membrane transporters.

The top upregulated genes include 2 TonB-receptors, a transcription factor, and genes related to amino acids and pyrimidine metabolism (Table 3).

**Table 3:**
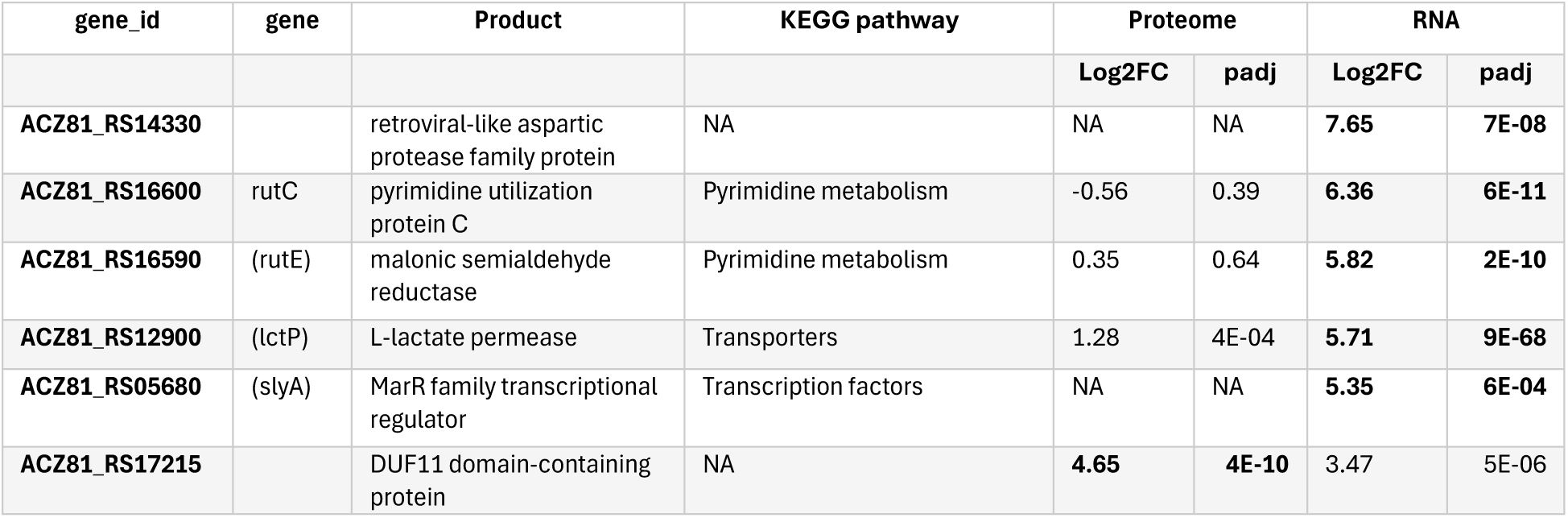
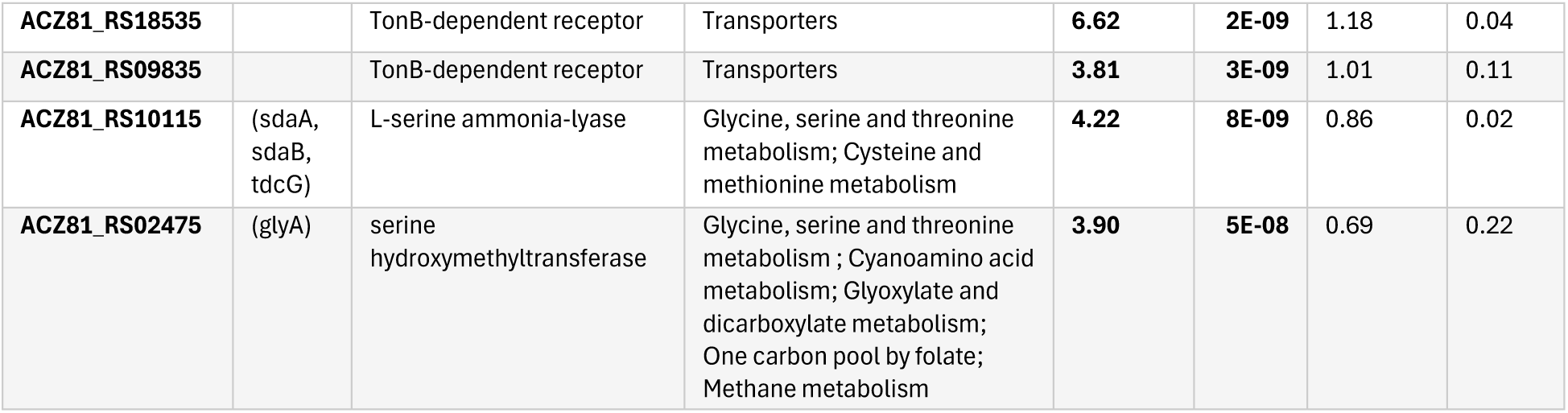
Top *Alteromonas* upregulated genes and proteins in long-term survival.

The most downregulated genes are related to the flagellar assembly, or their function is not known. All the five most downregulated transcripts are either upregulated in the proteome, or their proteome is not available (Table 4).

**Table 4:** Top *Alteromonas* downregulated genes and proteins in long-term survival.

| Gene id | gene | product | KEGG pathway | Proteome |  | RNA |  |
| --- | --- | --- | --- | --- | --- | --- | --- |
|  |  |  |  | Log2FC | padj | Log2FC | padj |
| ACZ81_RS07715 |  | MBL fold metallo-hydrolase | NA | 0.42 | 0.69 | -5.33 | 1E-13 |
| ACZ81_RS03025 |  | hypothetical protein | NA | NA | NA | -5.29 | 2E-21 |
| ACZ81_RS11430 |  | DUF1852 domain-containing protein | NA | NA | NA | -5.02 | 2E-17 |
| ACZ81_RS05055 | (fliC, hag) | flagellin | Flagellar assembly | 2.08 | 2E-04 | -4.99 | 8E-16 |
| ACZ81_RS04970 | (flgN) | flagellar protein FlgN | Flagellar assembly | NA | NA | -4.63 | 5E-96 |
| ACZ81_RS18760 |  | hypothetical protein | NA | -3.33 | 4E-04 | -1.80 | 7E-04 |
| ACZ81_RS04240 |  | HlyD family efflux transporter periplasmic adaptor subunit | Structural proteins | -3.49 | 3E-08 | -1.60 | 0.08 |
| ACZ81_RS10345 |  | c-type cytochrome | NA | -3.42 | 8E-09 | -1.50 | 0.10 |
| ACZ81_RS21255 |  | hypothetical protein | NA | -4.63 | 3E-07 | -1.42 | 0.02 |
| ACZ81_RS02160 |  | divergent polysaccharide deacetylase family protein | Function unknown | -3.24 | 6E-04 | 1.07 | 0.05 |

### GOSEQ enrichment in *Alteromonas*

During long term N starvation in co-culture with *Prochlorococcus*, the transcriptome and proteome results suggest that *Alteromonas* slows down some ecologically-relevant processes. This includes chemotaxis and motility and secretion. Surprisingly, ribosomes and translation are upregulated in both proteomes.

In the coculture, several metabolic pathways are upregulated, including metabolism of carbohydrates, amino acids and nucleotides, as well as nitrogen metabolism and transporters. These indicate a possible metabolic shift, where *Alteromonas* may feed on overflow photosynthate carbohydrates from *Prochlorococcus* (Calfee et al., 2022; Eigemann et al., 2022; Roth-Rosenberg et al., 2021).

**Fig 8:**
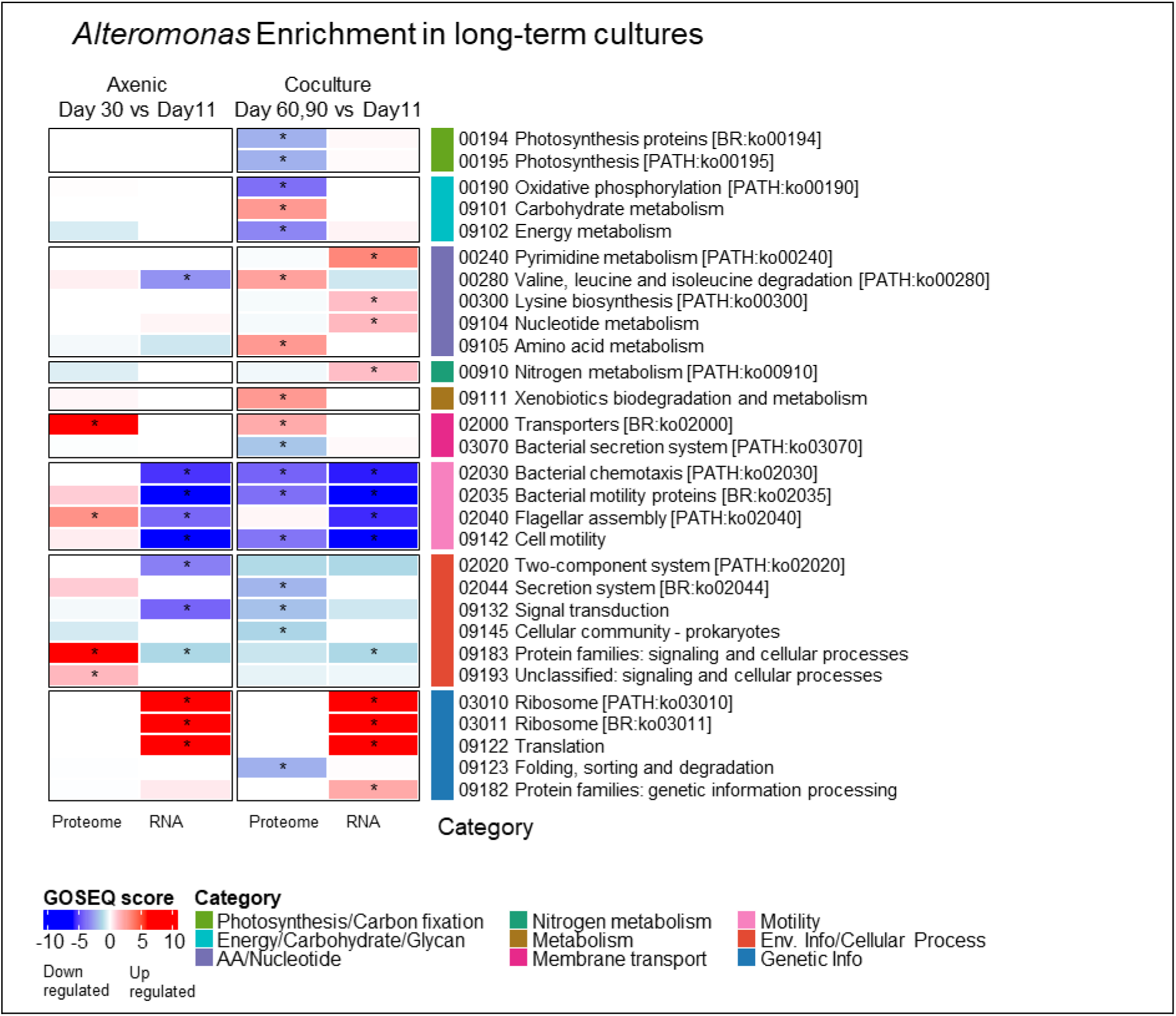
Enriched KEGG pathways in *Alteromonas* during Axenic death and long-tern survival in coculture. Pathways enriched when comparing death and long-term survival (days 60,90) to the initial timepoint.

## Discussion

The hypothesis that *Prochlorococcus* is nitrogen-limited is substantiated by the expression profiles, which demonstrate an upregulation of enzymes and transporters related to nitrogen metabolism and uptake. It is possible that, at the same time, *Prochlorococcus* avoids major changes to its C:N ratio by slowing down carbon fixation and reducing the production of nitrogen rich ribosomes (fig 3), (Domínguez-Martín et al., 2017; Szul et al., 2019; Tolonen et al., 2006). Notwithstanding this limitation, *Prochlorococcus* maintained viability throughout the prolonged starvation period, suggesting that a source of bioavailable nitrogen, albeit in minute quantities, remains accessible (fig 1). One potential nitrogen source is ammonium, as evidenced by the substantial production of ammonium in the axenic *Alteromonas* cultures (fig 1). These concentrations are not detected in the co-culture, a phenomenon that may be attributed to rapid turnover and immediate assimilation of all available ammonium by both co-culture members. Alternatively, it is possible that *Alteromonas* degrades some of the metabolites into smaller molecules that are bioavailable for *Prochlorococcus*, for example, it may degrade proteins into bioavailable amino acids.

Regarding *Alteromonas*, the observed reduction in motility may signify a phenotypic shift from a planktonic to an aggregated lifestyle. In terms of metabolism, upregulated carbohydrates pathways support the hypothesis that carbon is provided by *Prochlorococcus*, either by active exudates, or by recycling of dead cells (Grossowicz et al., 2017; Roth-Rosenberg et al., 2021; Weissberg et al., 2025). There is a complicated set of upregulated pathways related to amino acids and nucleotide metabolism, which is clearly driven by the coculture condition. Interestingly, we do not observe a metabolic shift in *Prochlorococcus*.

What constitutes the *Alteromonas* ‘helper’ phenotype? The prevailing hypothesis in the literature posits that a primary benefit involves the degradation of toxic reactive oxygen species (ROS) by *Alteromonas* (Ma et al., 2018; Morris et al., 2012). This mechanism likely plays a role in our experiment as *Alteromonas* upregulated ROS degradation enzymes in the coculture but not in the axenic controls (sup fig S2). Previous ROS degradation studies were performed in replete media (Ma et al., 2018). It appears that this mechanism remains active even when *Prochlorococcus* is subjected to nitrogen stress.

Remineralization by extracellular enzymes is another helper mechanism that have been suggested (Weissberg et al., 2025). These include extracellular proteases and aminases. *Alteromonas* upregulated extracellular proteolytic enzymes in both conditions, in cocultures as well as when grown axenically (sup fig S2). This is likely in response to nutrient stress as well as increasing levels of organic matter in the media, e.g., from decomposing dead cells. While degradation of proteins by these enzymes contributes to the recycling of nutrients in the media, their release does not seem to be in response to the presence of *Prochlorococcus*.

**Fig 9:**
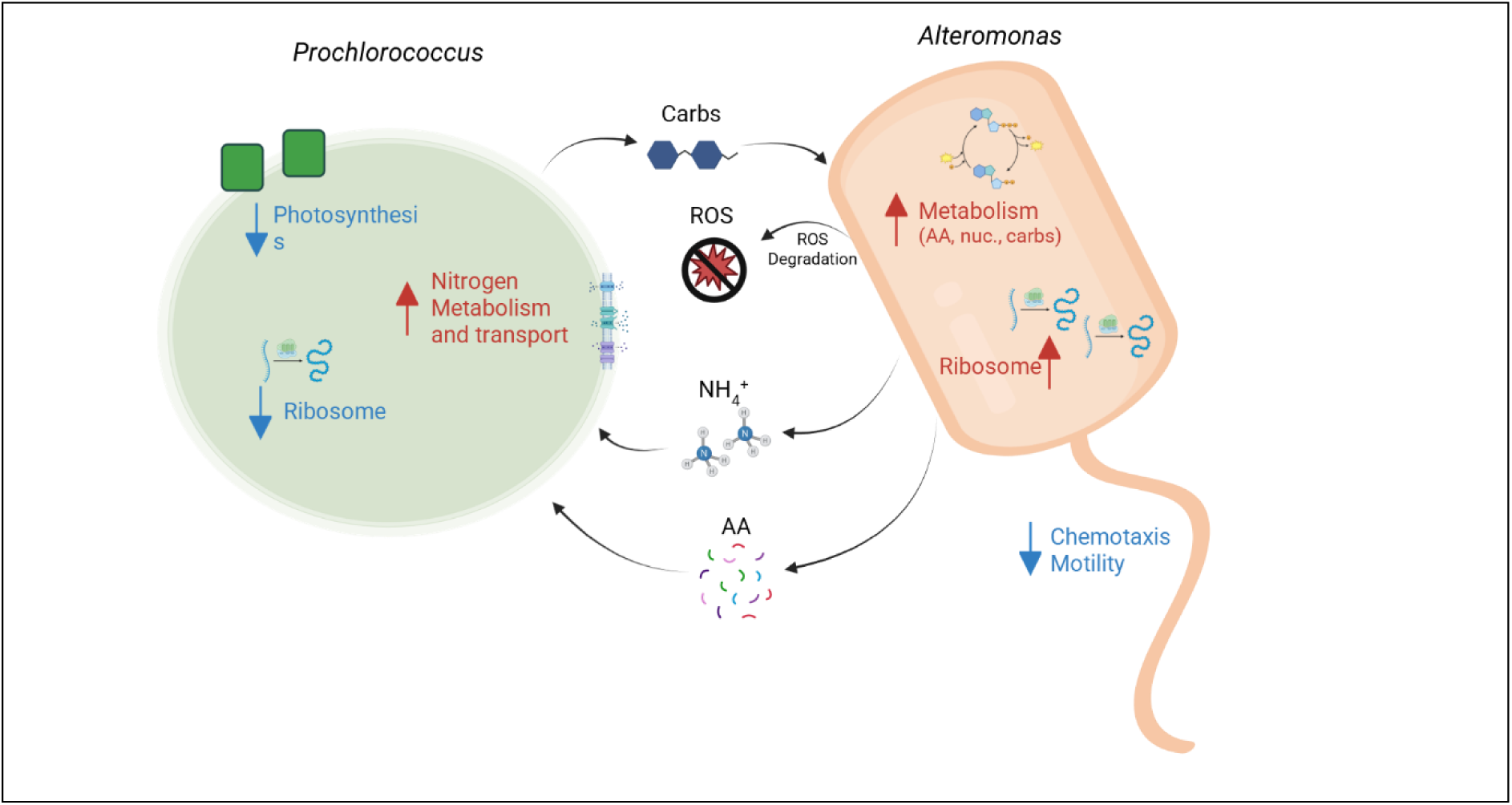
schematic illustration of most enriched pathways and interaction mechanisms in coculture during long term survival. AA: amino acids, nuc.: nucleotides, carbs: carbohydrates. ROS: reactive oxygen species. Created by biorender.com

There are several directions for continued exploration that we plan to address in a follow-up paper. What are the carbon and nitrogen nutrients being exchanged between *Prochlorococcus* and *Alteromonas*? While our study suggests a nutrient exchange, we did not identify the specific compounds exchanged. Previous studies suggest that *Prochlorococcus* may release compounds related to amino sugars, amino acids and purines (Coe et al., 2024; Kujawinski et al., 2023). None of these pathways is significantly enriched in our experiment. The answer may require a more intricate analysis, going beyond the level of pathways. We have conducted preliminary studies of the coculture using genome scale models (FBA – flux balance analysis, (Ofaim et al., 2021) but further studies are required. We also plan to run growth experiments, using amended media, to test which of the potential exchanges Alteromonas and Prochlorococcus can actually grow on. Another avenue to explore is to test which of the differentially expressed genes is important in the interaction and thus rank the affected pathways. We plan to do that using additional coculture experiments using a mutant library of a related *Alteromonas* strain (*Alteromonas* MIT1002).

What is the level of *Prochlorococcus* nitrogen limitation? Is the nitrogen stress lower in coculture due to availability of bioavailable nitrogen produced by *Alteromonas*? What are the expressions patterns of genes in the related pathways, and what can be deduced about their metabolic state? Related genes include nitrogen regulators (ntcA,..), nitrogen metabolism, nitrogen transporters, photosynthesis, HLIP, and ribosomes. We expect stronger response to increased nitrogen starvation, including increased upregulation of nitrogen related regulator and nitrogen metabolism and transport coupled with reduction in energy and translation.

Motility is strongly downregulated in *Alteromonas*. Does this indicate a shift to non-motile state? We plan to run motility assay to check. Reduced motility may be related to increased aggregation (Givati et al., 2023). In the case that *Prochlorococcus* and *Alteromonas* switch from free living to a more particle associated lifestyle, interactions between them are increased by their proximity. In that case, *Alteromonas*′ need for motility and chemotaxis will be reduced.

Finally, a large portion of the upregulated transcripts in *Prochlorococcus* are poorly characterized or their function is unknown. It would be interesting to analyze these. Are they related to response to environmental conditions in other studies? What is their location on the genome? Are they located on genomic islands? Mobile elements? Located in proximity to specific pathways or in operons? Recent advances in protein language models may also provide clues for the function of these genes. These models encode the nucleotide/amino acid sequence into an embedding. Often, genes with similar function may be close together in the embedding space.

## Conclusion

This study utilized a multi-omics approach, integrating transcriptomic and proteomic data, to dissect the physiological underpinnings of the synergistic interaction between the globally significant cyanobacterium *Prochlorococcus* and the heterotroph *Alteromonas* under conditions of prolonged nitrogen stress. Our findings demonstrate that the mutualistic association is critical for the long-term persistence of *Prochlorococcus* in the lab, allowing it to maintain viability long after its axenic counterpart has perished. The key mechanism enabling this extended survival is the nitrogen recycling function performed by *Alteromonas*. Evidence from our measurements indicates that *Alteromonas* effectively remineralizes organic nitrogen, primarily in the form of ammonium NH_4_^+^, which is then rapidly assimilated by both partners, suggesting a tightly coupled cross-feeding loop. The molecular data strongly support this conclusion. *Prochlorococcus* actively deploys nitrogen scavenging mechanisms through the upregulation of high-affinity transport and assimilation genes. Simultaneously, *Alteromonas* exhibits a phenotypic shift, likely associated with enhanced organic matter breakdown and reduced motility, which may facilitate a more substrate-attached or aggregated state optimized for localized nutrient remineralization. Collectively, these results extend the Black Queen Hypothesis by illustrating how the loss of core metabolic functions in a streamlined primary producer is compensated not only under oxidative stress but also through the continuous, active provisioning of essential nutrients, such as nitrogen, by a helper heterotroph when the external environment is exhausted. This dynamic, nutrient-releasing interaction is fundamental to understanding the resilience and stability of microbial communities in the vast, nutrient-poor regions of the ocean, providing crucial insights into the biogeochemical cycling that sustains global primary productivity.

## Methods

### Experiment setup

*Prochlorococcus* MED4 and *Alteromonas* HOT1A3 were grown in co-culture, as well as separately in axenic controls for 90 days. Cultures were grown triplicates of 750ml in 1L bottles using lowN-pro99 media (pro99 with NH_4_ ^+^ concentration reduced to 100μm instead of 800μm). Growth was measured continuously by measuring Fluorescence as well as cell counts using BD FACSCanto™ II Flow Cytometry Analyzer. Nutrients (NH_4_^+^, PO_4_) levels in the media were measured weekly. The viability of the heterotrophs was measured by MPN on day 89, and the viability of the *Prochlorococcus* in the co-cultures was measured by transfers to fresh media on days 31, 60, 89. Samples for Omics analysis (RNASEQ, proteomics, metabolomics) were taken on days 11, 18, 31, 60, 89, by filtering on 25mm GFF and flash freezing and stored in −80°C until analysis. For each culture, triplicate bottles were used for continuous growth measurement as well as for sampling on day 89. Additional triplicate bottles were used at each timepoint.

The axenic *Prochlorococcus* controls were repeated in a second experiment because the sampling on days 11, 18 did not match the timing of log exponential growth and decline. Instead, Omics measurements were taken on days 7, 14 to match med4 axenic growth dynamics.

### Fluorescence, Flow cytometry and MPN

Bulk chlorophyll fluorescence (FL) (ex440; em680) was measured every 3-5 days using a Fluorescence Spectrophotometer (Cary Eclipse, Varian). Samples for flow cytometry were taken every 3-5 days, fixed with glutaraldehyde (0.125% final concentration), incubated in the dark for 10 min and stored in −80 °C until analysis. 2 μm diameter fluorescent beads (Polysciences, Warminster, PA, USA) were added as an internal standard. Data was acquired and processed with FlowJo software. Flow cytometry was performed unstained to count *Prochlorococcus* cells followed by staining with SYBR Green I (Molecular Probes/ ThermoFisher) to count *Alteromonas* cells. Flow Cytometry results were processed by R CATALYST package.

MPN (most probable number) was calculated by transferring 20 ul of the culture in triplicate into a 96 well plate with marine broth and creating a 1:10 dilution series. OD (600nm) was measured on times 0 and after 24 hours of incubation in the dark. The probable number of *Alteromonas* viable cells is reflected by the dilution in which OD was higher after 24 hours.

### RNA SEQ

Samples for RNA collected on 25mm GF/F filters, preserved in 1 ml RNAlater and stored in −80. Total RNA was extracted from three biological replicates using the NucleoSpin RNA Stool kit (Macherey-Nagel) according to the manufacturer’s protocol with the following modifications. Filters thawed on ice, removed from the RNAlater and cut into 4 pieces (using ethanol clean scissors and forceps) and place in 15 ml tube with MN beads type A. 750 µl RST1 buffer and 200 µl NucleoZOL added to the filter and the tubes vortexed for 10 min. All next steps were according to the kit’s protocol. Total RNA eluted with 50 µl RNase free H_2_O. RNA quality and quantity were measured using Tape station and Qubit. The RNA quality, expressed as the RNA integrity number (RIN) was determined by electrophoresis using the 2200 TapeStation system (Agilent). RNA concentration was measured using Qubit 4 Fluorometer (Thermo Fisher). rRNA was removed using Illumina Ribo-Zero Plus rRNA Depletion Kit (Illumina) according to the manufacturer’s instructions, followed by generation of mRNA libraries using NEBNext® Ultra™ II Directional RNA Library Prep Kit for Illumina® (New England Biolabs) according to the manufacturer’s instructions. Sequencing libraries were sequenced on Illumina NextSeq 2000. Sample preparation and sequencing were conducted by the “Technion Genome Center,” Life Science and Engineering Interdisciplinary Research Center, Technion, Haifa, Israel.

Reads were trimmed using trimmomatic to remove TruSeq Adapter (TruSeq3-SE), quality checked using fastqc, rrna removed by sortmerna (silva-bac-16s-id90, silva-bac-23s-id98). reads were mapped by bwa mem to the genomes of MED4 (GCF_000011465.1) and HOT1A3 (GCF_001578515.1). Quality of the mapping was checked by qualimap. Common reads, defined as reads with high quality mapping to both genomes, were removed from both bam files using picard FilterSamReads. Reads were counted per genome using htseq-count with minimum quality 30.

### Proteomics

Samples for Proteins collected on 25mm GF/F filters, dried and stored in −80. Before the extraction, filters thawed on ice and 700ul Lysis buffer (40 mM EDTA, 50 mM Tris pH=8.3, 0.75 M Sucrose) added to the filter. Samples incubated with rotation at 37°C for 20 min. After incubation samples were sonicated in water bath for 10 min and centrifuge for 5 min at maximum speed on a table-top microfuge. The liquid transferred to a new 2 ml tube and centrifuge again. After second centrifugation the liquid transferred to a new Lobind protein Eppendorf (without the pellet) and the samples stored in −80. the BCA protein assay used to quantify total protein in the samples. The samples were digested with trypsin using the S-trap method. The resulting peptides were analyzed using nanoflow liquid chromatography (nanoAcquity) coupled to high resolution, high mass accuracy mass spectrometry (TIMS-TOF). Each sample was analyzed on the instrument separately in a random order in discovery mode.

Raw data was processed with FragePipe v17.1. The data was searched with the MSFragger search engine against your proteome databases appended with common lab protein contaminants and the following modifications: fixed carbamidomethylation on C, Oxidation on M and deamidation on NQ. Quantification was based on the MaxLFQ method, based on unique peptides.

### Analysis of Differential expressed genes and proteins

Differential gene expressions were analysed in R (4.2.0) by DESeq2 (2_1.38.2). Differential Protein abundance was analysed by DEqMS (1.16.0). KEGG pathways were identified by GhostKOALA. And pathway enrichment performed by GOSeq (1.50.0). Plotting by ggplot2 (3.4.1) and ComplexHeatmap (2.14.0).

## Code availability

The code used to analyse the data in this paper is available on github, https://github.com/wosnat/CC1A3.

## Acknowledgement

We thank Waseem Bashir for his assistance in the MPN assay and Tal Ben-Ezra for measuring NH_4_^+^ concentration. Shira Givati and Yara Soussan for good advice and assistance.

## Supplementary figures

**Sup fig S1:**
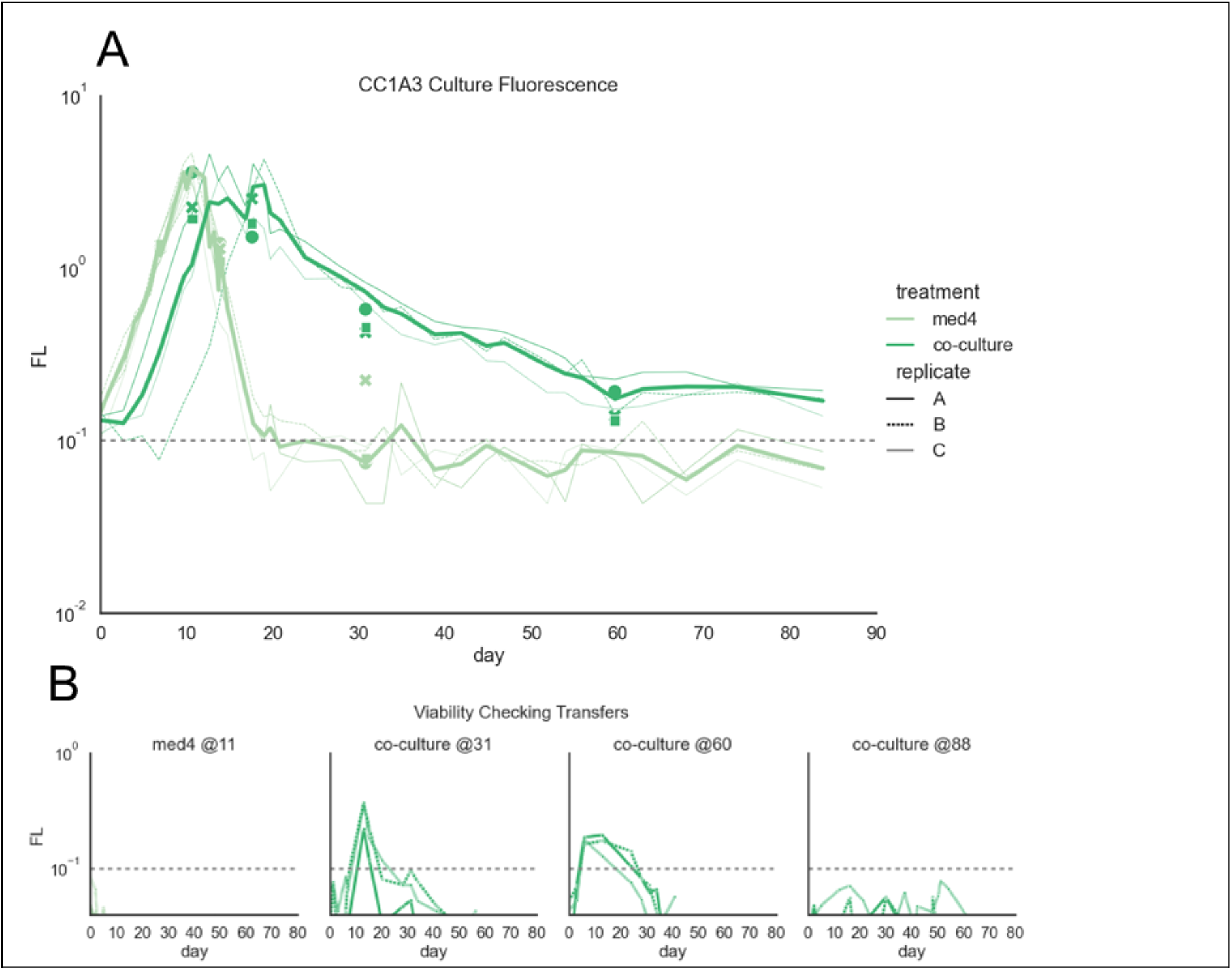
viability of the strains throughout the experiment. A. *Prochlorococcus* growth as measured by Fluorescence (med4 – axenic control, co-culture – *Prochlorococcus* in co-culture). B. Viability of *Prochlorococcus* as measured by transfer to fresh media.

**Sup fig S2:**
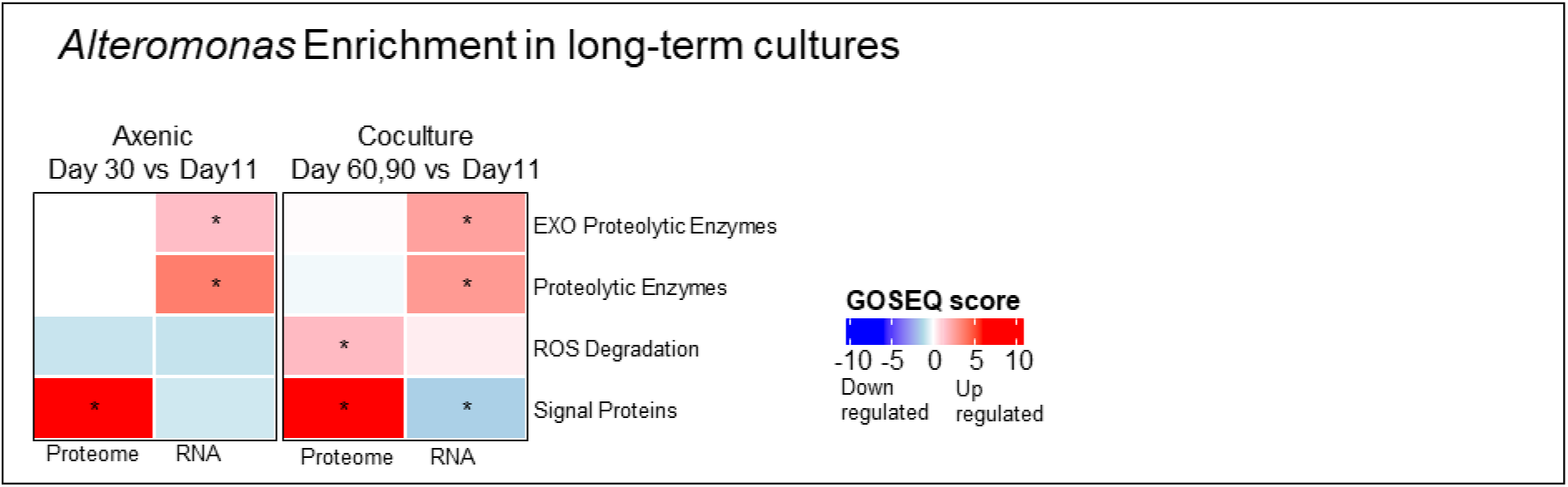
Enrichment of *Alteromonas* genes related to possible interaction mechanisms. Genes related to exoproteolytic enzymes and to degradation of toxic ROS.

